# Predicting the irrelevant: Neural effects of distractor predictability depend on load

**DOI:** 10.1101/2024.08.30.610431

**Authors:** Troby Ka-Yan Lui, Jonas Obleser, Malte Wöstmann

**Author notes:** **Corresponding author:** Malte Wöstmann, Department of Psychology, University of Lübeck, Lübeck, Germany, Maria-Goeppert Strasse 9a, 23562 Lübeck. **Data Accessibility** Pre-processed data supporting the results in this article are openly available: https://osf.io/m3pj2/.

## Abstract

Distraction is ubiquitous in human environments. Distracting input is often predictable, but we do not understand when or how humans can exploit this predictability. Here we ask whether predictable distractors are able to reduce uncertainty in updating the internal predictive model. We show that utilising a predictable distractor identity is not fully automatic but in part depends on available resources. In an auditory spatial n-back task, listeners (*n* = 33) attended to spoken numbers presented to one ear and detected repeating items. Distracting numbers presented to the other ear either followed a predictable (i.e., repetitive) sequence or were unpredictable. We used electroencephalography (EEG) to uncover neural responses to predictable versus unpredictable auditory distractors, as well as their dependence on perceptual and cognitive load. Neurally, pairs of targets and unpredictable distractors induced a sign-reversed lateralization of pre-stimulus alpha oscillations (∼10 Hz) and larger amplitude of the stimulus-evoked P2 event-related potential component. Under low versus high memory load, distractor predictability increased the magnitude of the frontal negativity component. Behaviourally, predictable distractors under low task demands (i.e., good signal-to-noise ratio and low memory load) made participants adopt a less biased response strategy. We conclude that predictable distractors decrease uncertainty and reduce the need for updating the internal predictive model. In turn, unpredictable distractors might mislead proactive spatial attention orientation, elicit larger neural responses, and put higher demand on memory.

## 1. Introduction

Depending on our goals, some signals in the environment are relevant targets and others are irrelevant distractors. Selective attention enhances mental representations of targets and suppresses distraction. Research in psychology and neuroscience has revealed sensory and higher-order features that modulate distractor processing, as well as associated neural enhancement and suppression mechanisms (Chelazzi et al., 2019; Geng, 2014; Noonan et al., 2018; van Moorselaar & Slagter, 2020; Wöstmann et al., 2022). Sensory signals are often coined by statistical regularities that allow prediction. Here, to investigate how the human mind exploits distractor predictability, we leverage neural responses preceding and following predictable vs. unpredictable distractors in the electroencephalogram (EEG). To test the hypothesis that predicting distraction depends on available cognitive resources, we systematically vary load in an auditory spatial attention task.

Acoustic distraction is inescapable since humans cannot easily “hear away” or “close their ears” to avoid irrelevant sound. Behavioural and electrophysiological investigations have shown that cues about auditory deviants reduce distraction and modulate stimulus-evoked responses (e.g., Horváth et al., 2011; Sussman et al., 2003). There is a rich literature on the Mismatch Negativity (MMN) response in the EEG, which shows that the human auditory system extracts stimulus relations in a sequence of (task-irrelevant) sounds to form expectations about upcoming events (e.g., Bendixen et al., 2012; Wacongne et al., 2012; Winkler et al., 2009). While such post-stimulus neural responses can provide insights on the encoding of (un)predictable sound and their integration with previously formed expectations, pre-stimulus neural signatures are though to signify how the neural system prepares for an upcoming stimulus that can or cannot be predicted (van Moorselaar et al., 2020). The power of neural alpha oscillations (∼10 Hz) is modulated both in anticipation of and following the presentation of competing auditory stimuli (Wöstmann et al., 2016). Alpha power modulation is sensitive to predictive benefits of acoustic input (Wilsch et al., 2015; Wöstmann et al., 2015), likely reflecting changes of neural enhancement versus suppression (Schneider et al., 2021). A comprehensive understanding of processing distractor predictability thus requires investigation of neural responses preceding and following distractors.

Putative distractor predictability effects are not expected to be fully automatic but instead to vary with levels of perceptual and cognitive load (Lavie, 2005). Molloy and colleagues (2019) found that processing of stochastic figure-ground patterns in task-irrelevant auditory stimuli decreases with higher visual perceptual load, suggesting a dependence on domain-general resources. If predicting distraction decreases under high load, this might have different implications. It could be that distractor predictability is not picked up under high load and preparatory processing is unnecessary. Alternatively, predictability might be used for preparatory processing, but high load prevents effective use of such preparation to modulate distractor processing. Here, we leverage pre-and post-distractor neural responses to distractors under varying levels of load to disentangle these alternative mechanisms.

Irrespective of task load, there are two competing views about processing distractor predictability, which will be contrasted in this study. First, predictability might increase saliency and enhance attention capture by the distractor (for an investigation challenging this view, see Southwell et al., 2017). This would increase neural responses to predictable distractors and potentially bias anticipatory attention to predictable distractors. Second, in line with predictive coding theory (Clark, 2013; Friston, 2010), predictable distractors might induce smaller prediction errors as the internal predictive model requires less updating. This would decrease neural responses to predictable distractors and potentially bias anticipatory attention to unpredictable rather than predictable distractors.

Benefits of spatial predictions (i.e., ‘predicting where’) and temporal predictions (i.e., ‘predicting when’) have been studied in some detail. Visual attention research has shown that the human brain is sensitive to statistical regularities regarding the spatial occurrence of distractors. If presented at one location with higher probability, distractors induce lower processing cost (B. Wang & Theeuwes, 2018) and fewer saccades are made to the high-probability distractor location (B. Wang et al., 2019). This suggests suppression of the high-probability distractor location. Temporally predictable (i.e., rhythmic) auditory distractors induce some benefits on target detection (Andreou et al., 2011; Makov & Zion Golumbic, 2020) and on secondary performance metrics (Lui & Wöstmann, 2022). However, identity predictions (i.e., ‘predicting what’; also referred to as ‘formal predictions’) are somewhat less well understood and induce dissociable auditory neural responses (Schwartze et al., 2013). Foreknowledge about the identity of an upcoming speech distractor reduces behavioural distraction cost (Röer et al., 2015). Furthermore, high-probability auditory distractors reduce interference with target detection (Daly & Pitt, 2021). These findings are compatible with the view that negative templates, which include distractor features, are employed for rejection (Arita et al., 2012). However, the associated neural mechanisms are at present largely unclear.

We here employ an auditory spatial attention paradigm wherein the temporal onset and the spatial position of an upcoming distractor are fully predictable, whereas its identity is either predictable or not. We hypothesize that predictable distractors reduce attention capture by the distractor and lower the neural processing demand, especially if perceptual and cognitive resources are available under low task load. Our findings support the notion that predictable distractors reduce the need for updating the internal predictive model, but processing distractor predictability is not fully automatic and depends instead on the availability of domain-general resources.

## 2. Methods

### 2.1 Participants

Thirty-three university students who were either native German speakers or non-native German speakers with high German language proficiency, participated in the EEG experiment for either course credits or €10/hour. One participant’s demographic information was lost but this participant’s data were included in all analyses. Participants (demographic information of remaining 32 participants: 25 females and 7 males, mean age = 24.25 years, SD = 3.89), provided written informed consent. According to self-report, they were right-handed (mean Edinburg Handedness Inventory score = 81.29; Oldfield, 1971), had normal hearing, and had no neurological or psychological disorders. All experimental procedures were approved by the local ethics committee of the University of Lübeck.

### 2.2 Stimuli and Procedure

Participants performed an auditory spatial n-back task (Fig. 1A) with manipulation of working memory load (1-vs. 2-back), perceptual load (target-to-distractor signal-to-noise ratio, SNR; 0 dB vs. –10 dB), and distractor predictability (predictable vs. unpredictable). Auditory stimuli were German numbers from 1 to 8, spoken by a female talker, and were shortened to 350 ms using Praat (version 6.1.16).

**Figure 1.**
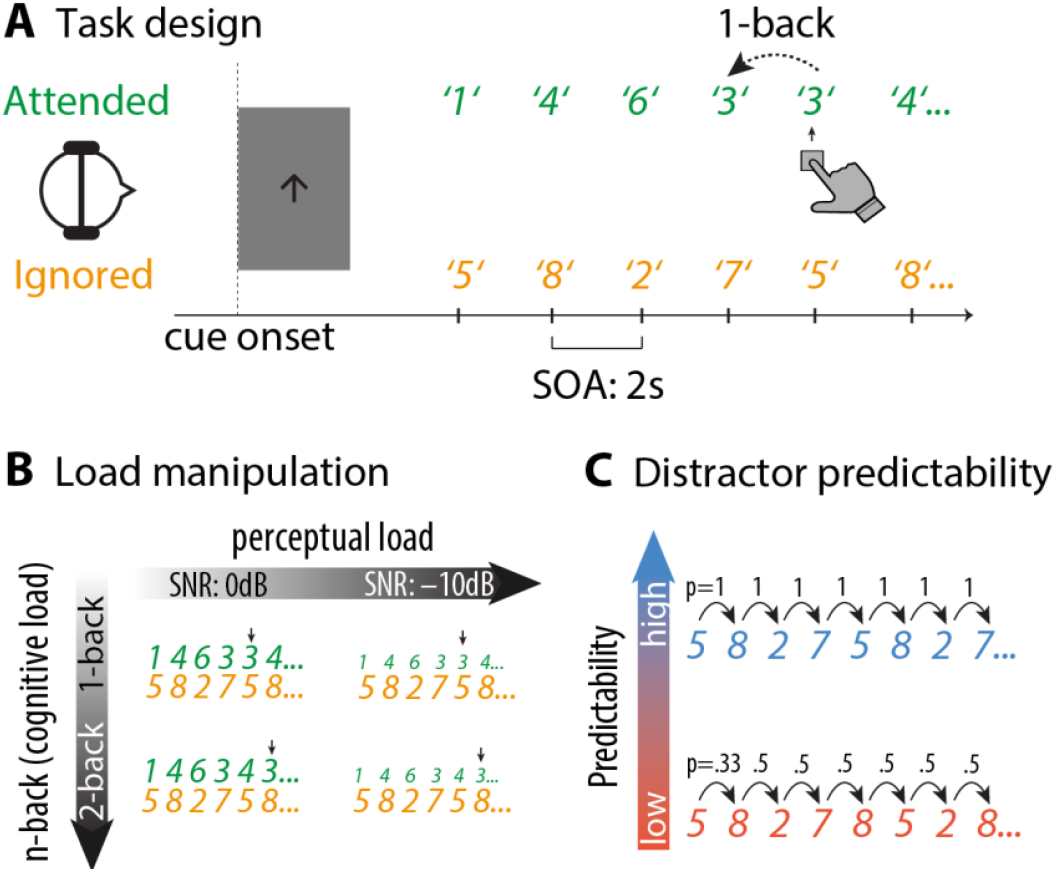
(**A**) Task design. Each block started with a spatial cue (arrow pointing left or right), indicating the to-be-attended side and the to-be-performed task (1-back: single-arrow, 2-back: double-arrow). Participants were presented with two competing streams of spoken numbers. They had the task of pressing a button when the current number on the to-be-attended side matched the previous number (in the 1-back condition) or the penultimate number (in the 2-back condition). (**B**) Task load was manipulated along two dimensions. First, the signal-to-noise ratio (SNR) of the target relative to the distractor stream was either 0 or –10 dB (visually displayed as differences in font size). Second, memory load was higher when participants performed a 2-back compared to a 1-back task. Arrows indicate putative button presses corresponding to the n-back task. (**C**) The distractor stream could either be predictable (blue; repeating in cycles of four numbers) or unpredictable (red). Numbers on arrows indicate transitional probabilities. In the unpredictable distractor stream, a given number was selected with the constraints that it was different from the previous and penultimate number, resulting in p = .33 for the second item (i.e., one number selected at random from three alternatives) and p = .5 for all subsequent numbers (i.e., one number selected from two alternatives).

Before each block, a visual cue (arrow) was presented in the centre of the screen to indicate the to-be-attended side (left or right) and the working memory load in that block (arrow with one line: 1-back, arrow with two lines: 2-back). Target numbers were presented monaurally on the cued side and distractor numbers were presented monaurally on the other side. Participants were instructed to attend to the cued side (target stream) and ignore the other side (distractor stream). For each trial, the target and the distractor numbers were presented simultaneously. Onset-to-onset interval between two numbers of a stream was 2s.

The working memory load was manipulated by the number of targets participants had to maintain in memory. In the target stream, a number sequence consisting of the target numbers was presented in pseudo-randomised order. Participants had to press the response button whenever the current target matched with the target 1 or 2 numbers prior to the current number in the 1-back and 2-back conditions, respectively. In each block, 20% of the presented numbers contained an n-back, where participants were supposed to press a button.

Perceptual load was manipulated as the signal-to-noise ratio (SNR) between the target and distractor sound intensities (Fig. 1B), which was analogous to the noise manipulation in a visual study of perceptual load (Gutteling et al., 2022). Specifically, the target stream was either presented at the same intensity as the distractor stream at ∼70 dB SPL or 10 dB SPL softer than the distractor stream. Participants were not informed about the SNR before each block. Instead, they were told prior to the main experiment that the stimulus intensity in the main experiment may vary from block to block.

Distractor predictability was operationalised as the transition probability of the distractor numbers in each block (Fig. 1C). In a predictable block, a randomly generated four-number pattern was presented repeatedly over the block, resulting in a transition probability of 1 for each distractor number. In an unpredictable block, the same four numbers were presented in a pseudo-randomised order, with the constraints that each number was different from the previous and penultimate number. The constraints were implemented to avoid unwanted potential confounds such as repetition suppression (Grill-Spector et al., 2006) or negative priming (Tipper, 1985). This resulted in a transition probability of 0.5 for each distractor number after the first two numbers in a block. Importantly, participants were not informed about the distractor predictability manipulation prior to the experiment.

There were 16 blocks in total, with each unique block (e.g., SNR 0 dB, 1-back, predictable distractors) repeating twice in the experiment. For each participant, the numbers 1 to 8 were randomly sorted into two groups. In half of the blocks, one group of numbers served as targets while the other group served as distractors, and vice versa in the other half of the blocks with the same conditions. Similarly, participants attended to the left side in half of the blocks and to the right side in the other half of the blocks. There were 120 target/distractor pairs per block and 1920 target/distractor pairs for the whole experiment. The auditory materials were presented via Sennheiser headphones (HD 25-1 II). A response box (The Black Box Toolkit) was used to collect behavioural responses. Stimuli were presented with Psychtoolbox (Brainard, 1997) for Matlab (MathWorks, Inc., Natick, USA).

### 2.3 Behavioural data analysis

To study how distractor predictability modulates behavioural performance under different load conditions, we used signal detection theory (SDT) to compute sensitivity (d’), and criterion (i.e., bias; c), as implemented in the Palamedes toolbox (Prins & Kingdom, 2018). The first (for 1-back block) or first two (for 2-back block) trials of each block were excluded in the behavioural analysis as there would be no previous number to be compared to. A hit was defined as a button press when the target number matched with the previous number (for 1-back) or the penultimate number (for 2-back) within the 2-s response window. A false alarm was defined as a button press when the current number did not match with the previous number (for 1-back) or the penultimate number (for 2-back).

Extreme hit or false alarm rates (0 or 1) were adjusted with a corrected value, which was computed by dividing 1 by 2 times the number of trials (Macmillan & Kaplan, 1985). A value of 0 was replaced by the corrected value, while a rate of 1 was adjusted by subtracting the corrected value from 1. We employed repeated-measures ANOVAs with factors SNR, working memory load, and distractor predictability to investigate effects on hit rate, sensitivity, and criterion. For criterion, post-hoc paired samples *t*-tests were used to contrast pairs of conditions.

### 2.4 EEG recording and pre-processing

The experiment was executed in a sound-attenuated and electrically shielded room. EEG data were recorded using 64 Ag/Ag-Cl electrodes (actiCHamp, Brain Products, München, Germany) with an online bandpass filter from direct current (DC) to 280 Hz. The sampling rate was 1000 Hz. TP9 (left mastoid) and FPz served as online reference and ground electrodes, respectively. For all participants, the impedances of the electrodes were kept below 20 kOhm.

The EEG data were pre-processed using Matlab R2018a (MathWorks, Inc., Natick, USA) and the Fieldtrip toolbox (Oostenveld et al., 2010). First, continuous data were filtered (high-pass filter: 0.1 Hz; low-pass filter: 100 Hz) and then segmented into epochs of 2 s (–1 to 1s; time-locked to the target/distractor pair onset). Then, an independent component analysis (ICA) was computed. Artefactual components related to eye blink and muscle activity were identified by visual inspection of components’ time courses, topographic maps, and frequency spectra and rejected. On average across participants, 30.21% of components were rejected (SD = 7.13%). Next, bad channels were visually identified and interpolated. Afterwards, trials containing absolute EEG amplitudes exceeding 160 μV were excluded. EEG epochs were re-referenced to the average of all electrodes. The first (for 1-back) or two (for 2-back) trials of each block, which were excluded in the behavioural analysis, were also excluded in the EEG analysis.

### 2.5 ERP analysis

The EEG epochs were first baseline corrected (–0.2 to 0 s) and re-referenced to the average of mastoid electrodes (i.e., TP9 and TP10). Then, EEG epochs were averaged to compute the event-related potential (ERP), for experimental conditions separately.

We studied the P2 and sustained negativity (SN) components in more detail. The P2 time window was determined by the mean amplitude around the peak of the grand average ERP waveform across all conditions and all participants. The positive peak in the grand average ERP waveform was at 185 ms and the time window used to extract P2 amplitude was selected from 160–210 ms. As the effect of the sustained negativity was robust and stable across a relatively long time interval, the ERP data 400–800 ms were averaged to obtain SN amplitude. The electrode with the maximum amplitude for each ERP component, as well as the two adjacent electrodes to the left and right, were used to calculate the ERP amplitude. As a result, the P2 amplitude was calculated using electrodes FC1, FCz, and FC2, while the sustained negativity was calculated using electrodes F1, Fz, and F2.

We examined the effects of perceptual load, working memory load, and distractor predictability on each ERP component using single-trial linear mixed-effects models (using the *fitlme* function in Matlab). For each trial, we averaged the amplitudes of the EEG data at the time windows and electrodes of interest. Then, we regressed the ERP amplitude on the main effects and interaction effects of the predictors and participant ID as a random intercept.

### 2.6 Analysis of alpha lateralization

Single-trial EEG data were decomposed into time-frequency representations via a fast Fourier transform (FFT) with a moving time window of 500 ms (Hanning taper). Complex Fourier coefficients were obtained from –.7 to .7 s (steps of 0.05 s) relative to target and distractor onset, and in a frequency range from 1 to 50 Hz in steps of 1 Hz.

The attentional modulation index (AMI) was calculated on absolute power to quantify spatial attention deployment, separately for different load and predictability conditions. First, trials belonging to respective attend-left or the attend-right conditions were averaged to increase the SNR (hence, no single-trial statistical analysis was carried out for alpha lateralization as opposed to the ERP analysis). Then, AMI was obtained according to Equation 1.

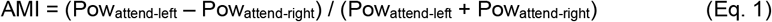

To test alpha lateralisation statistically, we averaged AMI across frequencies within the alpha band (8–12 Hz) and separately across a selection of posterior electrodes on the left and right hemisphere (TP9/10, TP7/8, CP5/6, CP3/4, CP1/ 2, P7/8, P5/6, P3/4, P1/2, PO7/8, PO3/4, and O1/2), which were employed in a previous study using the same EEG acquisition system (Wöstmann et al., 2019). For trials with high and low distractor predictability separately, paired samples t-tests comparing average AMI across left versus right electrodes were run across all time points, followed by false discovery rate (FDR) correction. The time windows that were significant after FDR correction (i.e., –0.5 to –0.35 s for unpredictable distractor condition; 0.3 to 0.5 s for both predictable and unpredictable distractor condition) were selected for further analysis.

In addition to AMI, we contrasted alpha power at electrodes ipsi-versus contralaterally relative to the focus of attention to obtain a time-resolved measure of alpha lateralization (Wöstmann et al., 2016). The alpha lateralisation index (ALI) was calculated according to Equation 2.

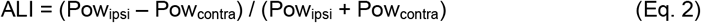

The ALIs in the selected time windows (i.e., T1: –0.5 to –0.35 s; T2: 0.3 to 0.5 s) were averaged for further statistical testing. A repeated-measures ANOVA was conducted to test the interactive effects of perceptual load (SNR), memory load (n-back), time window (T1 versus T2), and predictability (predictable versus unpredictable distractors) on ALI.

### 2.7 Effect sizes

For repeated-measures ANOVAs, we report partial eta-squared effect sizes (*η*_*p*_^*2*^). For t-tests, we report Cohen’s *d*. For mixed-effects models, we report the standardized partial effect size *r* for all relevant estimates, based on the *t*-value and the Sattherthwaite-approximated degrees of freedom (*df*) as shown in equation 3 (Carlson & Furr, 2009).

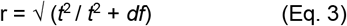

## 3. Results

The current study tested the effects of distractor predictability on the behavioural and neural dynamics of spatial attention, under varying levels of perceptual and cognitive load. Participants performed an auditory n-back task on a stream of target numbers presented via headphones to one ear (left or right), while a competing stream with predictable or unpredictable sequences of distracting numbers was presented to the other ear. Task load was manipulated along two dimensions: Perceptual load increased for low versus high target-to-distractor sound intensity ratios (–10 dB versus 0 dB SNR) and memory load increased for 2-back versus 1-back conditions.

### 3.1 Small effect of distractor predictability on response bias

Participants’ hit rates in the n-back task indicate good overall performance (Fig. 2A). Note that in n-back tasks of this kind, the number of items that are no n-back targets is high and only few of these non-targets are followed by a button press (i.e., small proportion of false alarms). Behavioural sensitivity (d’; Fig. 2B), which contrasts hit rate versus false alarm rate, is thus high. For completeness, we performed statistical analyses on both sensitivity and hit rate. Repeated-measures ANOVAs revealed better performance for 1-versus 2-back conditions (hit rate: *F*_*1,32*_ = 68.83, *p* < .001, *η*_*p*_^*2*^ = 0.683; sensitivity: *F*_*1,32*_ = 97.69, *p* < .001, *η*_*p*_^*2*^ = 0.753), but no main effects of perceptual load, predictability, or any interaction (all *p* > .13).

**Figure 2.**
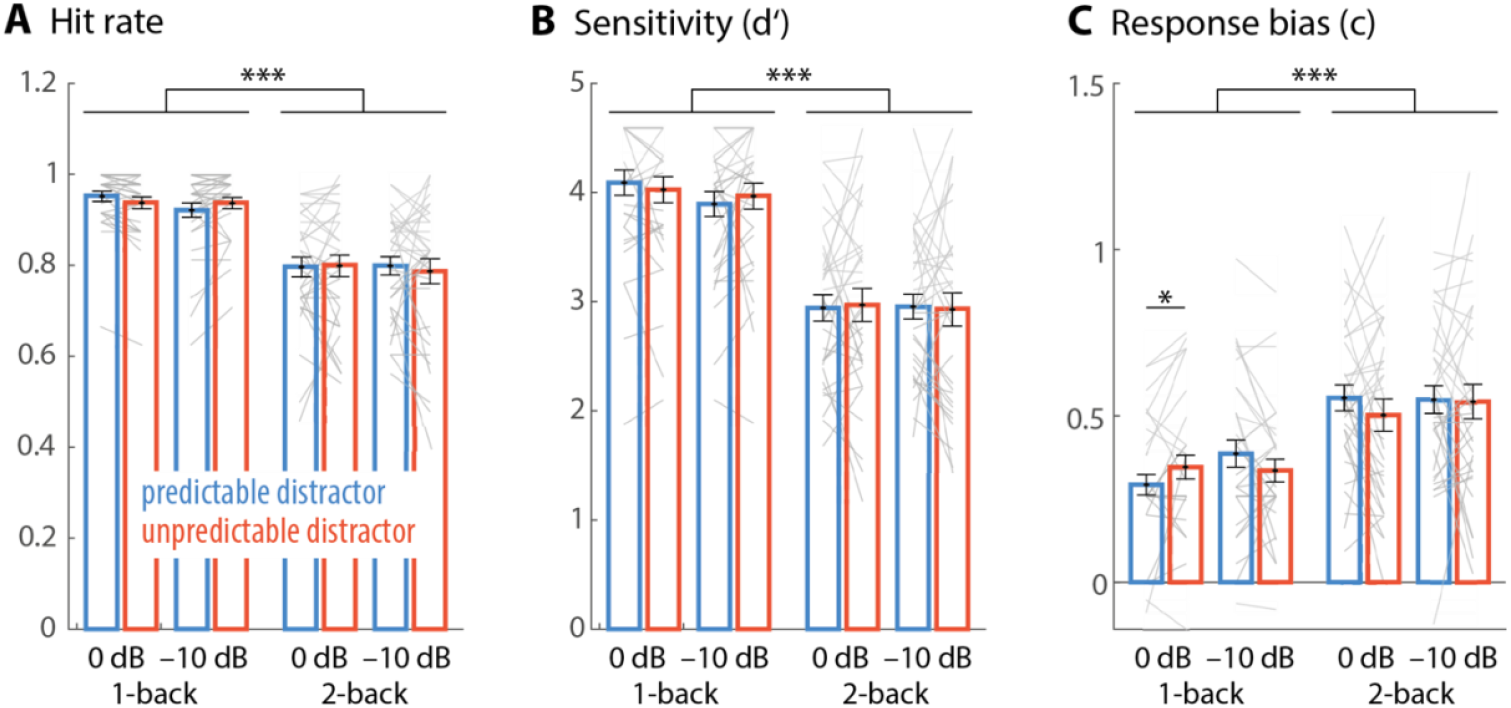
Bars and lines show average and single-subject hit rate (**A**), sensitivity (**B**), and response bias (**C**) for the different experimental conditions, respectively. Colour is coding for predictable (blue) versus (unpredictable) distractor sequences. Low load conditions were engendered by –10 dB SNR (low perceptual load) and 1-back (low memory load), respectively (i.e., most leftward bars). Error bars show ±1 SEM. * *p* < .05; *** *p* < .001.

Since response behaviour and metacognitive measures have recently been discussed to be sensitive to distraction effects (Kattner & Bryce, 2022; Lui & Wöstmann, 2022; Marsh et al., 2024), we analysed the effects of load and predictability on participants’ response bias. Most participants showed a conservative bias (Fig. 2C), which means that they tended to miss more

n-back targets than to report target-presence erroneously. Response bias was more conservative for the 2-versus 1-back condition (*F*_*1,32*_ = 30.85, *p* < .001, *η*_*p*_^*2*^ = 0.491). Furthermore, distractor predictability exhibited a small but statistically significant interactive effect with memory load and perceptual load on response bias (*F*_*1,32*_ = 4.19, *p* = .049, *η*_*p*_^*2*^ = 0.116). Response bias was closest to zero and relatively less conservative for predictable versus unpredictable distractors only in the easiest task condition with low perceptual load (i.e., 0 dB SNR) and low memory load (i.e., 1-back; *t*_*32*_ = –2.06; *p* = .048; *d* = 0.359).

### 3.2 Distractor predictability modulates neural dynamics of spatial attention

A major neural outcome measure to probe the dynamics of spatial attention deployment is the hemispheric lateralization of ∼10 Hz alpha oscillations. We found previously that the modulation of alpha lateralization over time relates to auditory spatial attention performance (Wöstmann et al., 2016) and is sensitive to explicit temporal cues (Wöstmann et al., 2021). Here, we tested whether lateralized alpha oscillations are sensitive to implicit manipulations of distractor predictability under varying levels of perceptual and cognitive load.

Figure 3 shows the Alpha Lateralization Index (ALI), which contrasts alpha power at parieto-occipital electrodes ipsi-versus contra-lateral to the focus of spatial attention. In general, a positive ALI reflects the typical pattern of higher ipsi-than contra-lateral alpha power during spatial attention. Time-windows of interest (T1 and T2) were selected by testing alpha lateralization against zero (see Methods for details). A repeated-measures ANOVA revealed a significant main effect of time window (*F*_*1,32*_ = 34.31, *p* < .001, *η*_*p*_^*2*^ = 0.52), indicating that ALI was more positive in the later (T2) compared to the earlier time window (T1). Furthermore, the main effect of SNR was significant (*F*_*1,32*_ = 4.60, *p* = .04, *η*_*p*_^*2*^ = 0.13), as well as the time window x SNR x predictability interaction (*F*_*1,32*_ = 4.61, *p* = .04, *η*_*p*_^*2*^ = 0.13), the predictability x time window interaction (*F*_*1,32*_ = 8.09, *p* = .008, η_p_^2^ = 0.2) and SNR x time window interaction (*F*_*1,32*_ = 4.65, *p* = .039, *η*_*p*_^*2*^ = 0.13).

**Figure 3.**
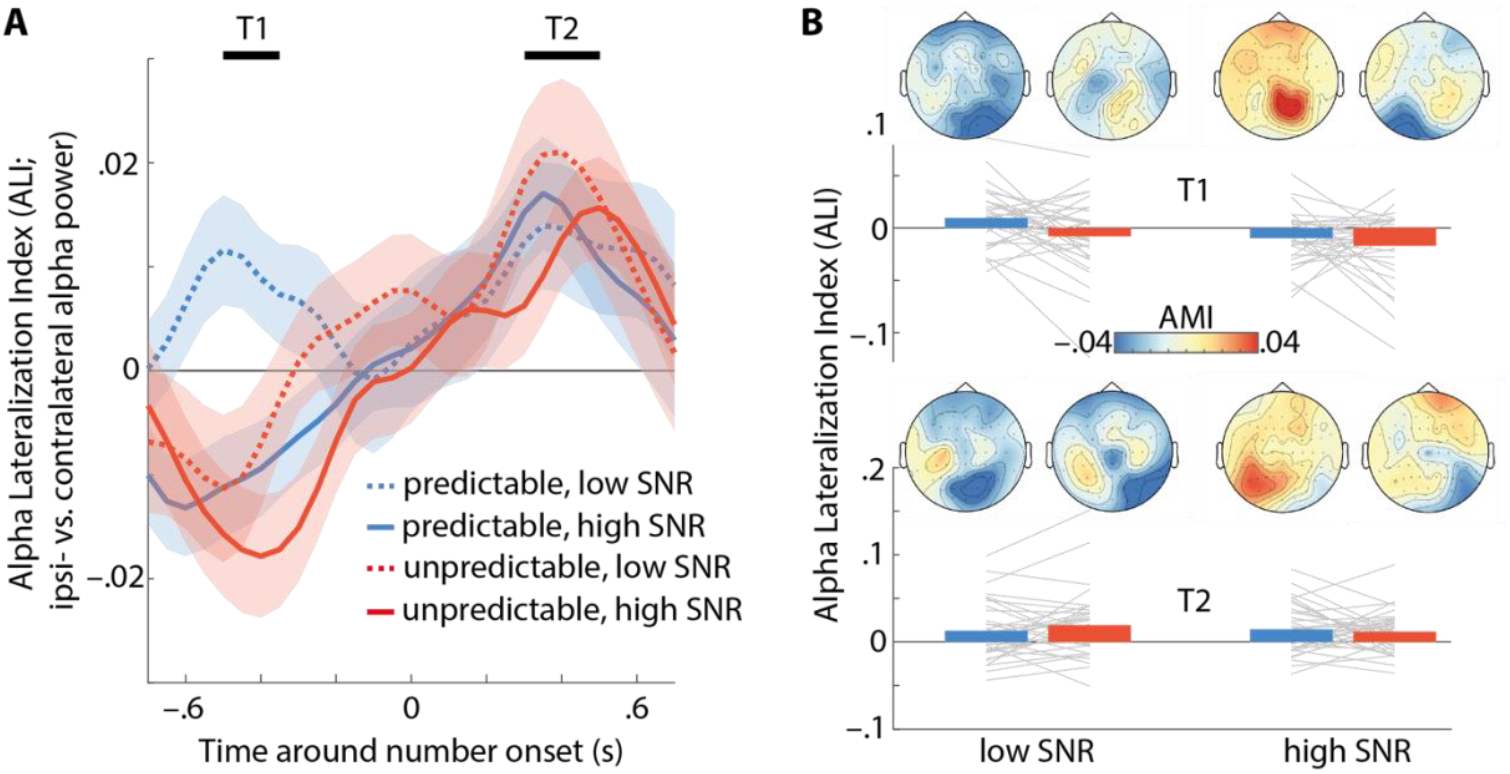
(**A**) Lines and shaded areas show the average Alpha Lateralization Index (ALI) ± 1SEM, respectively. Time windows T1 and T2 indicate pre-selected intervals wherein alpha lateralization differed from zero. (**B**) Bars and lines show average and single-subject ALI, respectively, for time windows T1 (top) and T2 (bottom), perceptual load conditions (low versus high SNR), and predictable (blue) versus unpredictable (red) distractors. Topographic maps show the alpha modulation index (AMI) for different conditions and time windows.

To resolve these interactions, we separated the data by time window. In the pre-stimulus time window (T1), there was a significant main effect of predictability (*F*_*1,32*_ = 7.55, *p* = .01, *η*_*p*_^*2*^ = 0.19), indicating a more negative ALI for unpredictable distractors. Moreover, the main effect SNR was significant (*F*_*1,32*_ = 9.83, *p* = .004, *η*_*p*_^*2*^ = 0.24), reflecting more negative ALI for higher SNR. All other main effects and interactions were not significant (all *p* > 0.18). In the post-stimulus time window (T2), there were no significant main effects or interactions of factors SNR, memory load, or predictability (all *p* > 0.14).

### 3.3 Independent and interactive effects of distractor predictability and load in the event-related potential

While distractor predictability effects on alpha power were found before stimulus onset (see above), we next investigated effects on the stimulus-evoked (i.e., post-stimulus) potential. The ERP showed the obligatory earlier P1, N1, and P2 components at central electrodes and a later negativity at frontal electrodes (Fig. 4A). The P2 component was independently affected by all three of our task manipulations: P2 amplitude was larger (i) when the target-to-distractor sound intensity ratio was higher (*t*_*61928*_ = 8.46, *p* < .001, *r* = 0.034), (ii) when memory load was higher in the 2-back compared with the 1-back condition (*t*_*61928*_ = 2.34, *p* = .02), and (iii) when the distractor was unpredictable (*t*_*61928*_ = 2.76, *p* = .006, *r* = 0.0111). No interactive effects of the three manipulations on P2 amplitude were found (all *p* > .24).

**Figure 4.**
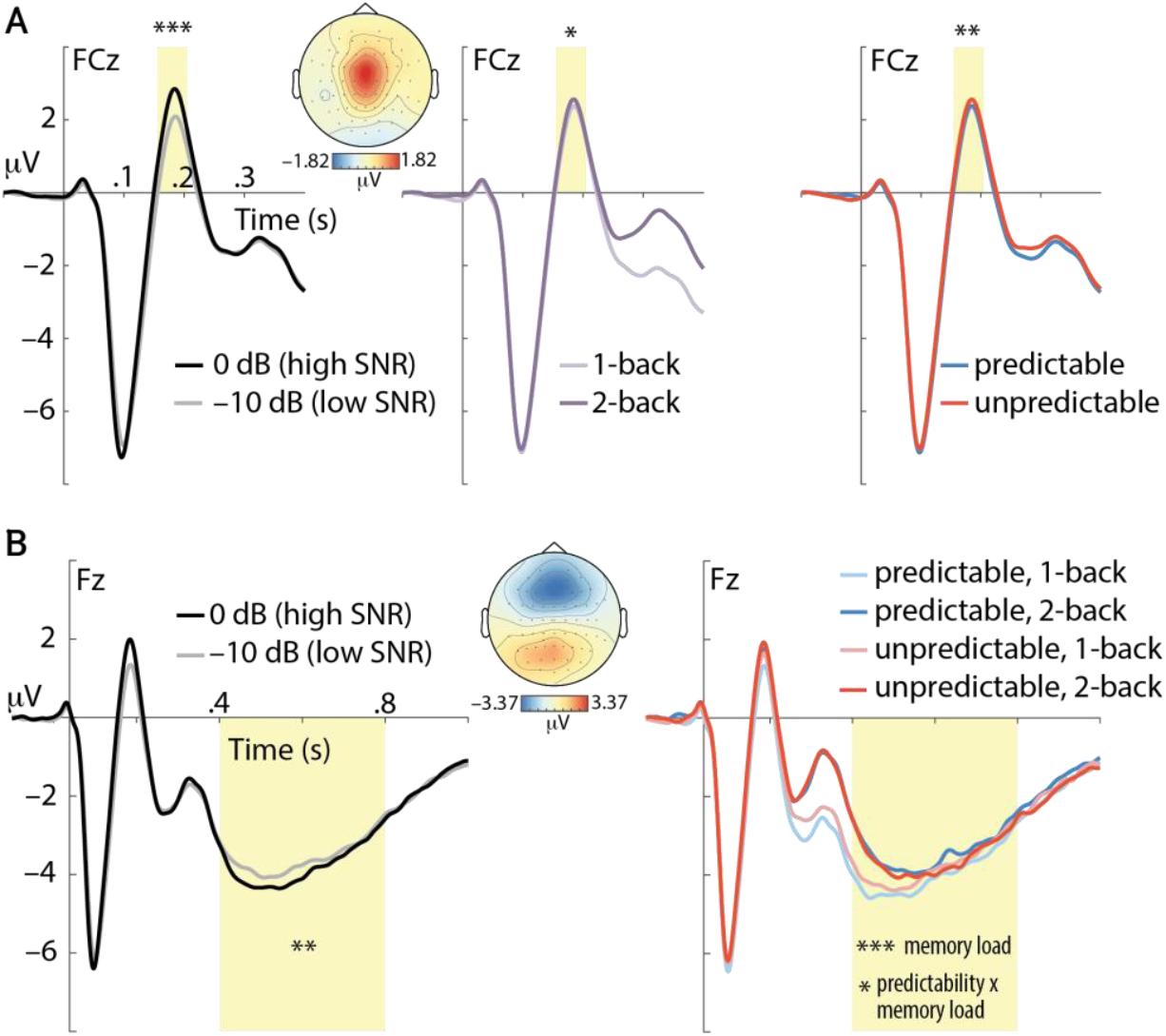
(**A**) Event-related potential (ERP) time-locked to the onset of pairs of numbers at electrode FCz. Main effects of perceptual load (left), memory load (middle), and distractor predictability (right) on amplitude in the P2 time window (highlighted in yellow) are shown. (**B**) The frontal negativity at electrode Fz (in time-window highlighted in yellow) was significantly modulated by perceptual load (SNR; left), memory load (right), as well as the memory load x predictability interaction. Topographic maps show average amplitude across all experimental conditions for the P2 (160– 210 ms) and frontal negativity (400–800 ms). * *p* < .05; ** *p* < .01; *** *p* < .001.

The frontal negativity is a long latency ERP component, which has previously been shown to relate to memory load in n-back tasks (e.g., Nowak et al., 2021). Here, we found that the magnitude of the frontal negativity increased (i.e., more negative amplitude) when SNR was high versus low (Fig. 4B; *t*_*61928*_ = 2.87, *p* = .004, *r* = 0.0115), as well as when memory load was low versus high (*t*_*61928*_ = 5, *p* < .001, *r* = 0.0201). Importantly, predictability modulated the frontal negativity interactively with memory load (*t*_*61928*_ = –2.25, *p* = .02, *r* = 0.009), meaning that predictability increased its magnitude when memory load was low (approaching statistical significance: *t*_*31076*_ = 1.83, *p* = .068, *r* = 0.0104) but the effect tended to reverse under high load (*t*_*30788*_ = –1.36, *p* = .174, *r* = 0.0078). No other interaction reached statistical significance (all *p* > .16).

## 4. Discussion

Does the human brain predict the identity of distracting input? And if so, how do available perceptual and cognitive resources constrain these predictions? To answer these questions, we employed an auditory spatial n-back task with competing streams of target and distractor items. Perceptual load increased with concomitantly lower SNR of target to distractor stimuli, and memory load increased for 2-back versus 1-back conditions. We investigated neural responses preceding and following predictable/expected distractor stimuli (for a similar approach, see Moorselaar, Daneshtalab, et al., 2021; van Moorselaar, Lampers, et al., 2021). While much previous research focused on spatial or temporal predictions, we investigated the predictability of distractor identity.

### 4.1 A spatial bias of attention to unpredictable distractors

The hemispheric lateralization of neural alpha oscillations (∼10 Hz) is a well-established signature of spatial attention across sensory modalities (auditory: Ahveninen et al., 2013; somatosensory: Haegens et al., 2011; visual: Worden et al., 2000). It has been suggested that supramodal alpha-band mechanisms in parietal cortical areas interact with sensory-specific control systems during spatial attention (Banerjee et al., 2011). Alpha power correlates negatively with neural activity assessed as the BOLD signal in functional magnetic resonance imaging (fMRI) in humans (Laufs et al., 2003) and with the firing rate of neurons in monkeys (Haegens, Nácher, et al., 2011). Thus, the implications of the contralateral decrease and ipsilateral increase in alpha power as respective reflections of target enhancement and distractor suppression during spatial attention have been discussed (Peylo et al., 2021; Schneider et al., 2021; van Moorselaar & Slagter, 2020). However, recent research questions the role of lateralized alpha oscillations for modulating neural gain measured as the stimulus-evoked response (Foster & Awh, 2019; Gundlach et al., 2020; Jensen, 2024; Morrow et al., 2023). Furthermore, EEG recordings alone can arguably not reveal the precise relation of oscillations and the computations underlying cognition and behaviour, which reflect in local neuron spiking (Snyder et al., 2015).

Although alpha lateralization is typically strongest in-between a spatial cue and stimulus onset (Wöstmann et al., 2019), we have shown post-stimulus modulation of alpha lateralization before, which eventually synchronized with the temporal structure of the acoustic input (Wöstmann et al., 2016, 2021). Here, we found two modulations of lateralized alpha power in pre- and post-stimulus time windows. First, higher ipsi-than contra-lateral alpha power approximately 400 ms after the onset of stimuli competing for spatial attention suggests spatial selection of target input and/or suppression of distraction (Wöstmann et al., 2016). Since this alpha modulation showed up after stimulus onset, it can be conceived as a neural signature of reactive attention deployment (Geng, 2014), potentially controlling the read-out of attended versus ignored sensory content. The absence of a distractor predictability effect on post-stimulus alpha lateralization suggests that reactive attention deployment happens irrespective of distractor predictability. Critically, however, the present study presented target and distractor items simultaneously and spatially confounded, that is, whenever the target was presented on the left, the distractor was presented on the right and vice versa. Thus, in contrast to previous investigations of ours (e.g., Orf et al., 2023; Wöstmann et al., 2019) the present design does not enable unambiguous association of neural responses with processing of targets versus distractors. Also, one might argue that probing participants’ responses to infrequent n-back items in the distractor stream might be a means to study attentional processing of the distractor more directly in behaviour. However, such a manipulation would arguably turn the distractor stream into an additional target, which is why it was avoided in the present study.

Second, alpha lateralization reversed in direction (i.e. relatively higher contra-versus ipsilateral alpha power) approximately 400 ms before the onset of pairs of targets and unpredictable versus predictable distractors (except for predictable distractors under high perceptual load, i.e., low SNR). We have previously shown that predictability of auditory target stimuli modulates alpha oscillations (Wöstmann et al., 2015) and that lateralized alpha power fluctuates during spatial attention and eventually goes back to baseline in-between stimulus presentation (Wöstmann et al., 2021).

The observed reversal of alpha lateralization might speak to spatial attention being biased to the location of the unpredictable distractor. This finding is reminiscent of the “ignoring paradox” (Moher & Egeth, 2012), which implies that distractor suppression is under some circumstances preceded by enhanced neural representation of the distractor (Donohue et al., 2018). Modulation of alpha lateralization accompanying involuntary shifts of spatial attention to task-irrelevant auditory deviants has been reported before (Weise et al., 2023). In theory, a bias of spatial attention to the location of an upcoming distractor can be considered sub-optimal, as it potentially interferes with the task goal. Possibly, spatial attention was biased to the distractor location to reduce uncertainty about unpredictable distractors.

In aggregate, higher contra-than ipsilateral alpha power for the unpredictable distractor condition suggests (i) that human listeners do extract statistical regularities in the temporal sequence of auditory distractors (Addleman & Jiang, 2019), and (ii) that listeners exhibit a spatial attention bias to the upcoming distractor in case it can be less well predicted.

### 4.2 Predictable distractors reduce neural processing demand

In the present study, the stimulus-evoked P2 component was sensitive to perceptual and cognitive load manipulations. Although P2 and N1 amplitudes are often correlated in empirical studies, it has been argued that the P2 can be conceived as an independent ERP component (Crowley & Colrain, 2004). Larger P2 amplitude for higher sound intensity (here: 0 dB vs –10 dB SNR) has been reported previously (Paiva et al., 2016) and likely reflects increased activity in auditory information processing. Similar to a previous investigation that found higher N1-P2 amplitude with higher memory load in an n-back task (Regenbogen et al., 2012), we found larger P2 amplitude for 2-back versus 1-back conditions.

Of high relevance for our research questions, P2 amplitude was suppressed for pairs of targets and predictable versus unpredictable distractors. P2 suppression has been associated with predictability of sound (Schröger et al., 2015). For instance, P2 amplitude was found to be suppressed for audio-visual versus audio-only speech presentation (van Wassenhove et al., 2005), as well as for temporally cued tones (Sowman et al., 2012). A further exploratory analysis of our data revealed smaller P2 amplitude for n-back targets versus non-targets in the present study (*t*_*61900*_ = –3.12; *p* = 0.0018, *r* = 0.0125), which agrees with the view that P2 suppression reflects a reduced prediction error. Lack of P2 suppression might be a signature of filling perceptual gaps (J. Wang et al., 2014), which explains larger P2 amplitude for pairs of targets and unpredictable distractor stimuli in the present study. Taken together, P2 suppression for pairs of targets and predictable distractors speaks to reduced prediction error (Knolle et al., 2013) and thus lower processing demand.

Starting approximately 400 ms after stimulus onset, a frontal negativity emerged in the ERP. This component is reminiscent of the sustained frontal/anterior negativity (SFN/SAN), which is a well-established signature of load during memory retention (e.g., Guimond et al., 2011; Lefebvre et al., 2013; Nolden et al., 2013). Sensitivity of the frontal negativity to memory has also been demonstrated in auditory n-back tasks, where its amplitude increased (Alain et al., 2009; Rämä et al., 2000) or decreased with higher load (Nowak et al., 2021). Given that we observed higher amplitude of the frontal negativity when the task demand was lower due to a higher SNR of acoustic stimuli or lower memory load, we presume that the frontal negativity reflects integration of the present stimulus with the existing memory trace, which would benefit from better acoustics and lower memory load. Further increased amplitude of the frontal negativity for pairs of targets and predictable distractors under low memory load (1-back) might thus indicate a distractor predictability-induced memory processing benefit in the present task.

### 4.3 Predicting distraction depends on available resources

Here, we manipulated perceptual and memory load (i) to test whether distractor predictability processing is automatic (i.e., independent of load) or contingent on limitations of available resources, and (ii) to explore whether distractor predictability effects are modulated stronger by perceptual load (presumably favoring early attentional selection; Lavie, 2005) or memory load (presumably favoring late selection; Zhang et al., 2006).

Distractor predictability effects in the present study were partly independent of load (pre-stimulus alpha and P2 modulation) and partly larger under low load: Response bias was least conservative for predictable distractors when perceptual and memory load were low, and the distractor-predictability increase in the frontal negativity ERP component was larger under low memory load. These effects suggest that processing of distractor predictability is not fully automatic but requires resources that are not available in case of high load. Constraints of available cognitive resources for predictive language processing have been reported before (Ito et al., 2018; Ryskin & Nieuwland, 2023). The present findings might also explain the absence of distractor predictability effects in some of our own previous studies where memory load was high in all conditions (Lui & Wöstmann, 2022; Wöstmann & Obleser, 2016). Mechanistically, it appears that if resources are available to exploit distractor predictability (i.e., under low load), memory integration of target items is facilitated.

Since distractor predictability effects in the present study interacted with perceptual and memory load, our results do not have strong implications about the exact type of resources necessary for distractor prediction. It has been debated to what extent perceptual load theory is applicable to attention in the auditory modality (Murphy et al., 2017) and whether SNR manipulations induce the same effect on perceptual load as other possible manipulations (such as increasing the number of items or target-distractor similarity). Of note, the high perceptual load condition in the present study (with target intensity lowered relative to distractor intensity) used an overall reduced sound intensity, which might have counteracted potential effects. Our results suggest that the dependence of distractor predictability processing on resources is gradual rather than discrete (i.e. “all or nothing”). That is, while effects of predictability on memory integration (reflected by the frontal negativity in the ERP) and response bias were modulated by load, spatial attention bias (reflected by pre-stimulus alpha modulation) and suppression of the stimulus-evoked P2 were independent of load.

One important design feature of the present study was that the temporal occurrence and spatial location of all stimuli were fully predictable in all experimental conditions. In this sense, distractor stimuli were highly predictable overall, whereas our main experimental manipulation only changed predictability along one dimension: distractor identity. Thus, one might speculate that distractor predictability effects would be larger in case of higher perceptual uncertainty about distractors in a context wherein the temporal occurrence and spatial position would vary unpredictably.

## 5. Conclusion

The present study shows that the listening brain does extract subtle statistical regularities from a sequence of irrelevant speech items. Prediction of distractors is not fully automatic but depends on the availability of perceptual and cognitive resources. As pre-stimulus oscillatory and post-stimulus evoked neural responses show, unpredictable distractors are more potent in misleading proactive spatial attention allocation and do increase subsequent distractor-processing costs. These findings help understand the potential benefits of predictable distractors for goal-directed neural processing and its dependence on perceptual and cognitive resource limitations.

## List of abbreviations

EEG: Electroencephalography
SNR: Signal-to-Noise Ratio
MMN: Mismatch Negativity
FFT: Fast Fourier Transform
AMI: Alpha Modulation Index
ALI: Alpha Lateralization Index

## Acknowledgments

This work was supported by Deutsche Forschungsgemeinschaft [grant number WO 2371/1– 1, to MW]. We thank two anonymous reviewers for their helpful comments on a previous version of this manuscript.

